# Transcriptional landscape of PTEN loss in primary prostate cancer

**DOI:** 10.1101/2020.10.08.332049

**Authors:** Eddie Luidy Imada, Diego Fernando Sanchez, Wikum Dinalankara, Thiago Vidotto, Ericka M Ebot, Svitlana Tyekucheva, Gloria Regina Franco, Lorelei Mucci, Massimo Loda, Edward M Schaeffer, Tamara Lotan, Luigi Marchionni

## Abstract

PTEN is the most frequently lost tumor suppressor in primary prostate cancer (PCa) and its loss is associated with aggressive disease. However, the transcriptional changes associated with PTEN loss in PCa have not been described in detail. Here, we applied a meta-analysis approach, leveraging two large PCa cohorts with experimentally validated PTEN and ERG status, to derive a transcriptomic signature of *PTEN* loss, while also accounting for potential confounders due to *ERG* rearrangements. Strikingly, the signature indicates a strong activation of both innate and adaptive immune systems upon *PTEN* loss, as well as an expected activation of cell-cycle genes. Moreover, we made use of our recently developed FC-R2 expression atlas to expand this signature to include many non-coding RNAs recently annotated by the FANTOM consortium. With this resource, we analyzed the TCGA-PRAD cohort, creating a comprehensive transcriptomic landscape of *PTEN* loss in PCa that comprises both the coding and an extensive non-coding counterpart.

## Introduction

Previous molecular studies have explored the genomic heterogeneity of prostate adenocarcinomas (PCa) revealing distinct molecular subsets characterized by common genome alterations (1–3). Among these molecular alterations, loss of the tumor suppressor gene phosphatase and tensin homolog *(PTEN)* – which is implicated in the negative-regulation of the PI3K-AKT-mTOR pathway – has been identified as one of the most common genomic drivers of primary PCa (4,5). Since alterations in the PI3K pathway are present in more than 30% of human cancers, the identification of an expression signature associated with *PTEN* loss has been investigated in different tumor contexts, including breast, bladder, lung, and PCa (6,7).

Assessment of *PTEN* status by fluorescence in situ hybridization (FISH) and immunohistochemistry (IHC) in large clinical PCa cohorts has shown a consistent association with adverse pathological features such as high Gleason score, extra-prostatic extension, as well as prognostic value for biochemical recurrence and cancer-related death (4,8). IHC-based assessment of *PTEN* status has been shown to correlate tightly with genomic alterations of the *PTEN* locus and captures not only loss of the gene, but also mutation and epigenetic changes that lead to *PTEN* functional inactivation(4,9,10) and the potential clinical utility of PTEN IHC as a valuable prognostic marker has been demonstrated previously (11–14).

Though PTEN is involved in a myriad of cellular processes spanning cellular proliferation to tumor microenvironment interactions (5), the transcriptional landscape related to *PTEN* expression has not yet been explored in depth, and the role of long non-coding RNAs (lncRNAs) remains elusive (15). These observations, added to the evidence that subtle *PTEN* downregulation can lead to cancer susceptibility (16), demonstrate the important role of *PTEN* in cancer biology but also highlight the need for additional studies.

Similarly, gene rearrangements of the ETS transcription factor, *ERG*, with the androgen-regulated gene Transmembrane Serine Protease 2 (*TMPRSS2*) are present in ~50% of PCa from patients of European descent. *TMPRSS2-ERG* fusion (herein denoted as *ERG*^+^ for fusion present and *ERG*^−^ for absence of fusion) has been shown to activate the PI3K-kinase pathway similarly to *PTEN* loss (17), leading to increased proliferation and invasion. Importantly, tumors harboring *TMPRSS2-ERG* rearrangements show an enrichment for *PTEN* loss (17,18). The co-occurrence of these two genomic alterations makes it challenging to dissect the contributions of each to the transcriptomic landscape.

The goal of this study was to elucidate the transcriptional landscape of *PTEN* loss in PCa through the analysis of two large and very well clinically-curated cohorts, for which *PTEN* and *ERG* status was assessed by clinical-grade IHC: The Natural History (NH) cohort, in which patients that underwent radical prostatectomy for clinically localized PCa did not receive neoadjuvant therapy or adjuvant hormonal therapy prior to documented distant metastases (19); and the Health Professionals Follow-up Study (HPFS) cohort in which the patients were followed for over 25 years (20). Based on IHC-assessed *PTEN* status for these cohorts, we built a *PTEN*-loss signature highly concordant across the independent datasets, in both presence and absence of *TMPRSS2-ERG* fusion. Overall, this *PTEN*-loss signature was associated with cellular processes associated with aggressive tumor behavior (*e.g*., increased motility and proliferation) and, surprisingly, with increases in gene sets related to the immune response. In addition, through our recently developed FANTOM-CAT/recount2 (FC-R2) resource (21) and copy-number-variation data, we expanded this signature beyond coding genes and report the non-coding RNA repertory resulting from *PTEN* loss.

## Methods

### Data collection and Immunostaining

All expression data used in this work were gathered from public domain databases. In this work, we made use of three cohorts: FC-R2 TCGA, Natural History (NH), and Health Professionals Follow-up Study (HPFS). Information about each cohort is summarized in Table 1. Information about *PTEN* status by immunohistochemistry for the HPFS cohort was readily available and therefore obtained from the public domain. For NH cohort samples, IHC staining for *PTEN* and *ERG* were performed using a previously validated protocol (22). Last, for TCGA we used the Copy Number Variation (CNV) called by the GISTIC algorithm to define *PTEN* status and expectation-maximization algorithm to define ERG status.

**Table 1.** Cohorts summary Table shows cohorts summary for the 3 cohorts used in this study: TCGA (only primary tumor samples with high Gistic scores were used); Health Professional Follow-up Study (all); and Natural History cohort (samples with IHC call available). PTEN-null represents samples with PTEN deletion and PTEN-intact regular primary tumors.

| Cohort | PTEN-null | PTEN-intact | N |
| --- | --- | --- | --- |
| TCGA | 95 | 321 | 416 |
| HPFS | 91 | 299 | 390 |
| Natural History | 56 | 151 | 207 |
| Total | 242 | 771 | 1,013 |

### Meta-analysis of NH and HPFS cohorts

We performed a meta-analysis approach using a Bayesian hierarchical multi-level model (BHM) for cross-study detection of differential gene expression implemented in the Bioconductor package XDE (23) on microarray-based cohorts to obtain a *PTEN*-null signature from *PTEN* IHC validated samples. The model was fitted using the delta gp model with empirical starting values and 1000 bootstraps were performed. All remaining parameters were set to default values. This analysis was also performed stratifying the samples by *ERG* status to evaluate the impact of the *ERG* rearrangement in the signature.

### Differential expression analysis in the TCGA cohort

A generalized linear model (GLM) approach coupled with empirical Bayes moderation of standard errors and voom precision weigths (24,25) was used to detect differentially expressed genes in the TCGA cohort. The models were adjusted for surrogate variables with the SVA package (26). Adjusted p-values controlling for multiple hypothesis testing were performed using the Benjamini-Hochberg method and genes with false discovery rate (FDR) equal or less than 0.1 were reported (27).

### Gene set enrichment analysis (GSEA)

The results from the meta-analysis performed in the NH and HPFS cohort were ranked by the weighted size effect (average of the posterior probability of concordant differential expression multiplied by the Bayesian effect size of each cohort). The results from the TCGA cohort were ranked by t-statistics. Ranked lists were tested for gene set enrichment. Gene set enrichment analysis (GSEA) was performed using a Monte Carlo adaptive multilevel splitting approach, implemented in the fgsea (28) package. A collection of gene sets (Hallmarks, REACTOME, and GO Biological Processes) were obtained from the Broad Institute MSigDB database. The androgen response gene set was obtained from Scheaffer et al (29). Gene sets with less than 15 and more than 1500 genes were removed from the analysis, except for the GO biological processes whose max size was set to 300 to avoid overly generic gene sets. The enriched pathways were collapsed to maintain only independent ones using the function collapsePathways from fgsea.

## Results

### Meta-analysis of Natural History and Health Professionals Follow-Up Study cohorts

We sought to obtain a consensus signature of *PTEN* loss that could be reproduced across independent cohorts. We utilized a meta-analysis approach leveraging a multi-level model for cross-study detection of differential gene expression (DGE). We fitted a Bayesian hierarchical model (BHM) for analysis of differential expression across multiple studies that allowed us to aggregate data from two previously described tissue microarray-based cohorts where *PTEN* and *ERG* status was determined by IHC (Table 1 and Figure 1) and we derived a *PTEN*-loss signature (Figure 2). In this analysis, we observed 813 genes for which the differential expression was highly concordant (Bayesian Effect Size (BES) ≥ 1, posterior probability of concordant differential expression (PPCDE) ≥ 0.95) (Table S1).

**Figure 1.**
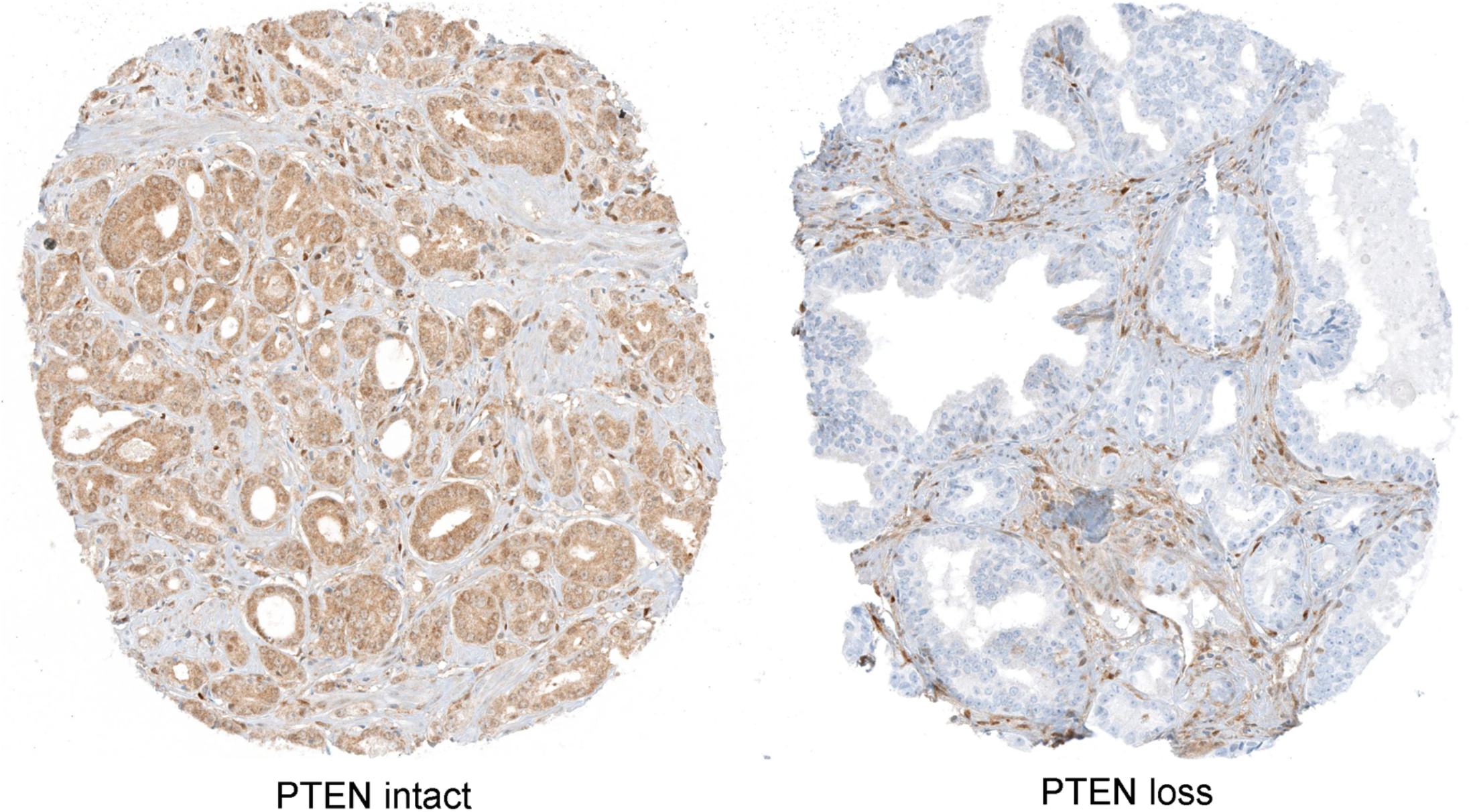
PTEN immunostaining in tissue microarray (TMA) spots from the Natural History Cohort. Left panel: intact PTEN protein is present in all sampled tumor glands (brown chromogen). Right panel: PTEN loss in all sampled tumor glands. Images reduced from 40X.

**Figure 2.**
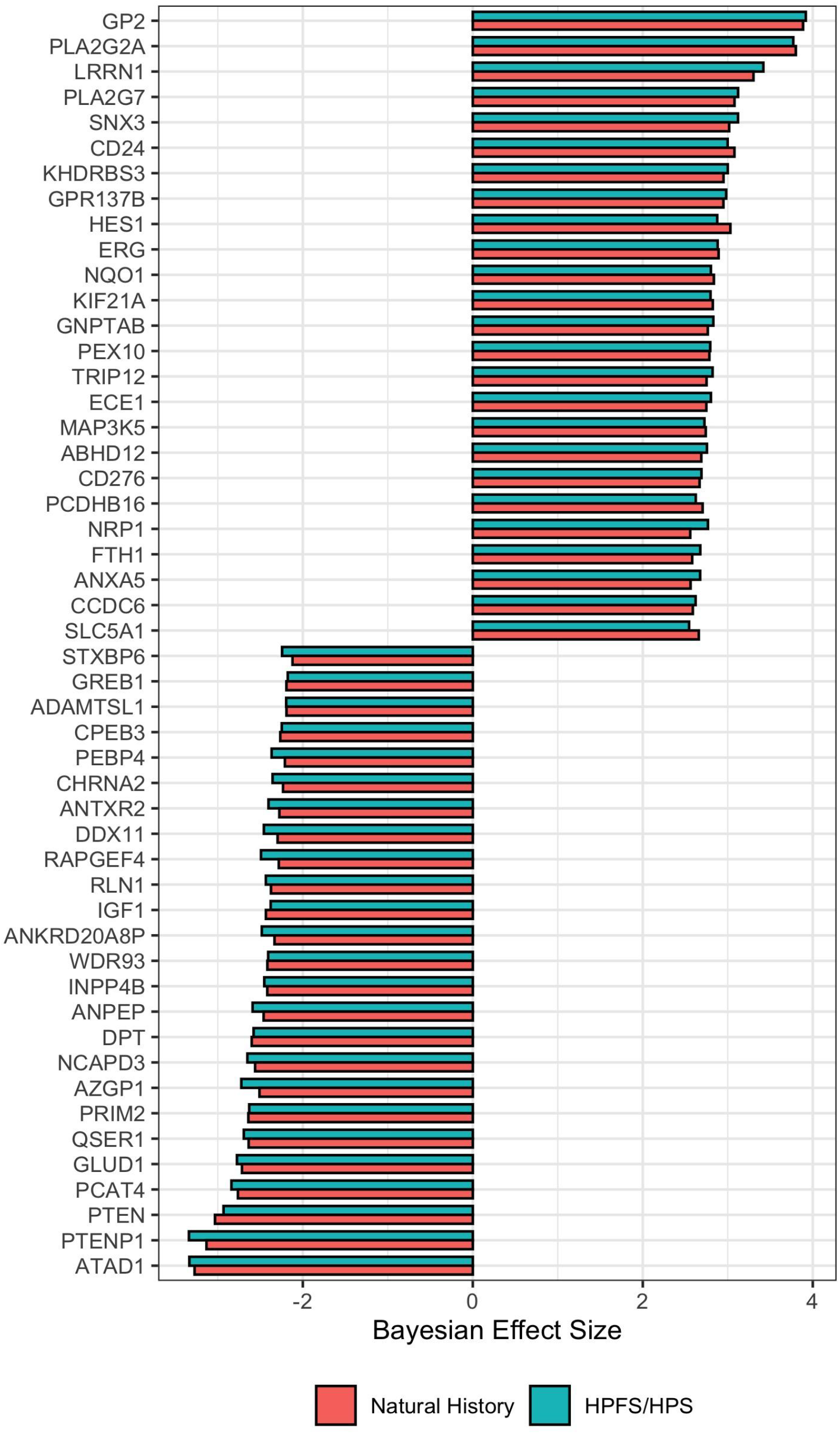
Cross-study meta-analysis of differential gene expression. Genes in the same loci as PTEN such as RLN1 and ATAD1 were found down-regulated. PTEN-null vs PTEN-intact meta-analysis of HPFS/PHS and NH cohorts with Bayesian Hierarchical Model for DGE using XDE showing the top 25 most concordant differentially up- and down-regulated genes. PTEN status were based on IHC assays.

The consequences of *PTEN* loss on cell cycle regulation and tumor cell invasion has been extensively reported previously (4,30,31). Accordingly, beyond *PTEN* itself, the top DEG genes in our signature reflected this profile (Figure 2 and Table S1). Dermatopontin (*DPT*) (BES = −2.59, PPCDE = 1) and Alanyl membrane aminopeptidase (*ANPEP*) (BES = −2.53, PPCDE = 1) were found down-regulated upon *PTEN* loss. Leucine-Rich Repeat Neuronal 1 (*LRRN1*) was among the genes up-regulated upon *PTEN* loss (BES = 3.36, PPCDE = 1). These and other genes found differentially expressed upon *PTEN* loss have all been shown to be associated with a more aggressive phenotype in several cancer types (5).

Notably, we found *ERG* among the top upregulated genes in the signature (Figure 2). As expected (18,32,33), *ERG* rearrangement was more common among cases with *PTEN* loss compared to intact *PTEN* in all cohorts (Fisher exact test, p ≤ 0.001). Given this enrichment, it was not surprising that *ERG* was among the most up-regulated genes in the BHM signature, as well as *PLA2G7*, which has been shown to be among the most highly overexpressed genes in *ERG*-rearranged PCa compared to those lacking *ERG* rearrangements (34). The presence of ERG and ERG-regulated transcripts in the *PTEN*-loss signature suggested that this signature might be confounded by enrichment of *ERG* rearranged tumors among the tumors with PTEN loss.

Since *ERG* rearrangements represent a major driver event in PCa and *PTEN* loss is enriched in *ERG*-rearranged tumors, we next investigated the role of ERG in our PTEN-loss signature. To this end, we repeated the Bayesian hierarchical model for the analysis of differential expression by stratifying the samples by *ERG* status. In the background with *ERG* rearrangement, we observed a similar signature to the previous overall *PTEN*-loss signature, but without the aforementioned *ERG*-associated genes (Supplementary figure S1 and Supplementary table S2). However, in the absence of *ERG* rearrangement, we could not find any significant differences between samples with or without *PTEN* loss. This was unexpected given that *PTEN* is a powerful tumor suppressor capable of triggering multiple molecular changes.

### Extending the PTEN-loss signature

To validate our PTEN loss signatures in an orthogonal cohort, we next examined the TCGA PRAD cohort (35), where *PTEN* status was estimated by genomic copy number (CN) assessment, which was closely aligned with PTEN gene expression (Figure S3). We recently developed a comprehensive expression atlas based on the FANTOM-CAT annotations. This meta-assembly is currently the broadest collection of the human transcriptome (21,36). These gene models include many novel lncRNA categories such as enhancers and promoters, allowing the signature to be further expanded beyond the coding repertoire. We used TCGA expression data from the FC-R2 expression atlas (21) to perform DGE analysis stratified by the *PTEN* status as derived from CN analysis. We also performed the same analysis in a stratified manner as in the HPFS and NH cohorts, using the ERG expression with expectation maximization (EM) algorithm to define ERG status given the bimodal nature of ERG expression in PCa. Interestingly, we were able to detect differential expression between *PTEN*-null and PTEN-intact samples without *ERG* rearrangement in the TCGA cohort, which used high-throughput sequencing as opposed to gene expression microarrays, suggesting that there the lack of signal in the previous analysis can be a reflection of the potential limitations with the later technology.

We observed 521 differentially expressed genes (DEG) when comparing *PTEN*-null and *PTEN*-wild-type samples (FDR ≤ 0.01, LogFC ≥ 1), of which 257 were coding genes and 264 were non-coding genes (Supplementary Table S3). When stratifying the samples by ERG status, we obtained 435 and 364 DEG in the background with and without ERG rearrangement (Supplementary Table S4 and S5), respectively, with similar proportions of coding and non-coding genes. Using Correspondence-at-the-top (CAT) analysis of the coding genes, we observed a higher concordance than expected by chance between the TCGA PTEN-loss signature and that from the BHM (Figure S4). This confirmed that CN is a reasonable proxy to IHC-staining in TCGA which allowed us to expand this signature beyond coding RNAs.

In this analysis, we were able to detect a variety of lncRNAs that are already known to be involved in PCa development and progression. Notably, several differentially expressed lncRNAs were already reported to be associated with PCa (37–46) (e.g. *PCA3, PCGEM1, SCHLAP1, KRTAP5-AS1, Mir-596)* (Supplementary Table S3-S5). *PCA3* is a prostate-specific lncRNA overexpressed in PCa tissue. Similarly, lncRNA *PCGEM1* expression is increased and highly specific in PCa where it promotes cell growth and it has been associated with high-risk PCa patients (41,42). On the other hand, *KRTAP5-AS1* expression has not been directly associated with PCa.

Also ranked high among lncRNAs differentially expressed were the lncRNAs *SChLAP1* and its uncharacterized antisense neighbor *AC009478.1. SchLAP1* is overexpressed in a subset of PCa where it antagonizes the tumor-suppressive function of the SWI/SNF complex and can independently predict poor outcomes (45,46). On the other hand, the role of *AC009478.1* in PCa development is still unknown. Interestingly, *SchLAP1* and *AC009478.1* expression is strongly correlated in the TCGA datasets only in PCa (R = 0.94, p < 2.2e-26) and bladder cancer (R = 0.85, p < 2.2e-26) (Figure S5).

Strikingly, a substantial proportion of lncRNAs associated with *PTEN* loss were not yet associated with PCa. Out of the 264 DE non-coding genes, 134 were novel and annotated only in the FANTOM-CAT meta-assembly annotation (Table 2). Among the FANTOM-CAT exclusive genes, those with the highest fold change in close proximity with coding genes were *CATG00000038715, CATG00000079217*, and *CATG00000117664* (Figure S6). These genes were mostly expressed in PCa as opposed to other cancer types in the TCGA dataset (Figure 3).

**Table 2.** Summary of differentially expressed genes between PTEN-null and PTEN-intact with logFC ≥ 1 and FDR ≤ 0.01 across different ERG backgrounds. Number in parenthesis shows the number of genes exclusive to the FANTOM-CAT annotations.

|  | PTEN-null vs PTEN-intact overall | PTEN-null vs PTEN-intact in ERG+ | PTEN-null vs PTEN-intact in ERG- |
| --- | --- | --- | --- |
| Coding genes | 257 (13) | 226 (7) | 185 (10) |
| Non-coding genes | 264 (134) | 209 (117) | 179 (82) |
| Total | 521 (137) | 435 (124) | 364 (92) |

**Figure 3.**
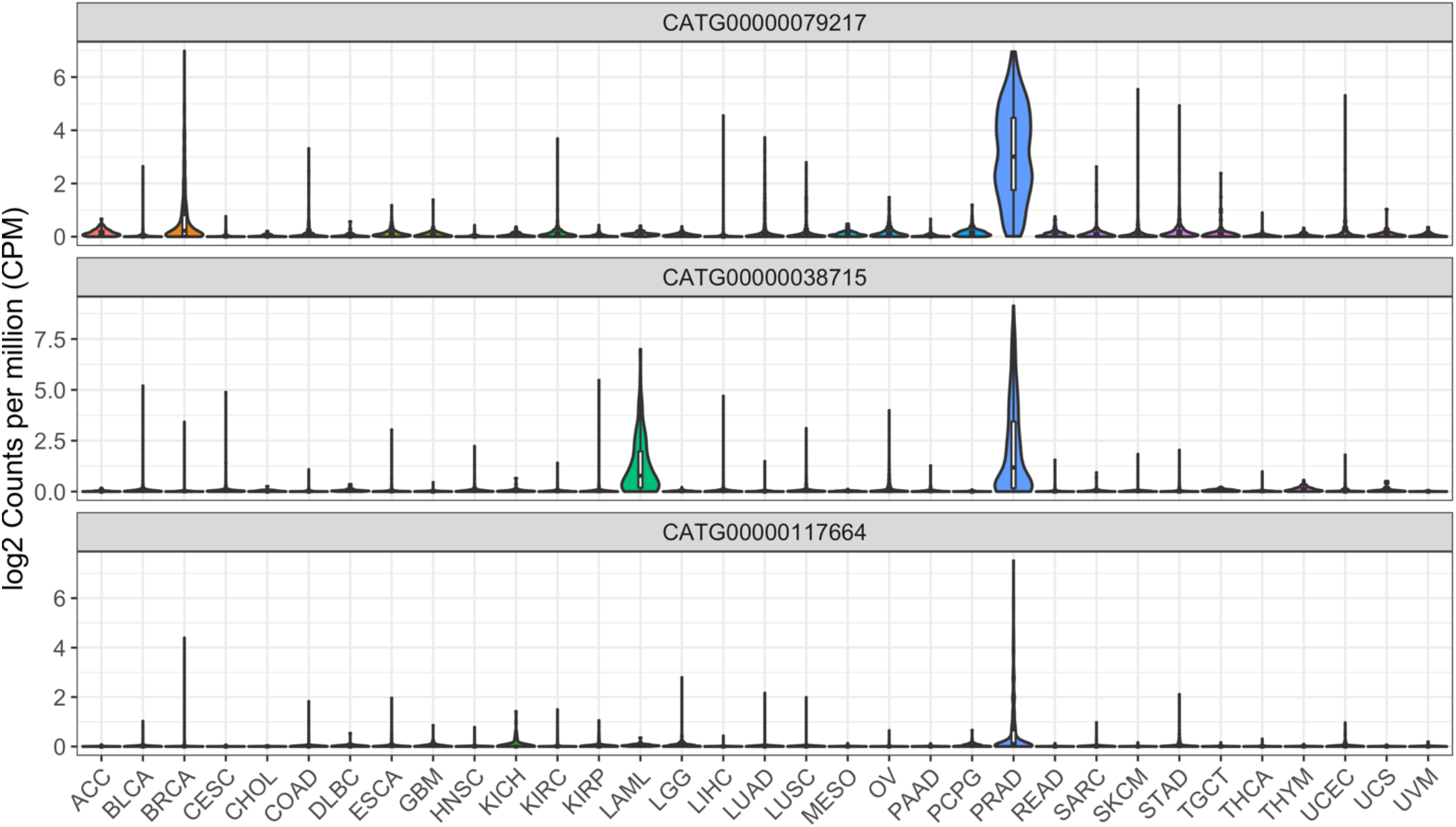
Expression profiles of novel FANTOM-CAT genes CATG00000038715, CATG00000079217 and CATG00000117664 across 33 cancer types. Violin-plots shows expression (log2 CPM+1) distribution.

Among the downregulated genes were *CATG00000038715* and *CATG00000079217. CATG00000038715* is in close proximity to *CYP4F2* and *CYP4F11*, encoding members of the cytochrome P450 enzyme superfamily. Expression of *CATG00000038715* and *CYP4F2* are highly correlated (R=0.91, p < 2.2e-16) in PCa, and expression of the former was highly specific for PCa (Figure S7). *CATG00000079217* is in close proximity to the coding gene *FBXL7*, an F-box gene which is a component of the E3 ubiquitin ligase complex. While expression of *FBXL7* and *CATG00000079217* showed only a weak correlation (R=0.14, p < 7.4e-4), *CATG00000079217* expression was notably higher in PCa and breast cancer than in other cancers, and it was moderately correlated with several PCa biomarkers (e.g. *KLK2, KLK3, STEAP2, PCGEM1, SLC45A3*) (41,42,47–51) (R=0.37-0.57, p < 2.2e-16) in TCGA.

*CATG00000117664* was among the most upregulated lncRNA and it is located near *GPR158*, a G protein coupled receptor highly expressed in brain. The expression between *GPR158* were correlated (R=0.54, p < 2.2e-16), and *CATG00000117664* expression was shown to be highly specific to PCa (52) (Figure S7).

### PTEN loss induces the innate and adaptive immune system

We performed Gene Set Enrichment Analysis (GSEA) using fgsea (28) and tested both the BHM- and TCGA-generated molecular signatures for enrichment in three collections of the Molecular Signature Database (MSigDB) (53,54): HALLMARKS, REACTOME, and GO Biological Processes (BP). Results were similar in both signatures, with positive enrichment of proliferation and cell cycle-related gene sets (e.g. MYC1 targets, MTORC1 signaling, cell cycle checkpoints, and DNA repair) and both innate and adaptive immune system associated gene sets (e.g. Neutrophil degranulation, MHC antigen presentation, interferon-alpha, and gamma) (Figure 4-5 and Supplementary Table S6-S20). The positive enrichment of MHC antigen presentation, interferon-alpha and -gamma in PTEN-null tumors is consistent with our previous study showing that the absolute density of T-cells is increased in PCa with PTEN loss (55).

**Figure 4.**
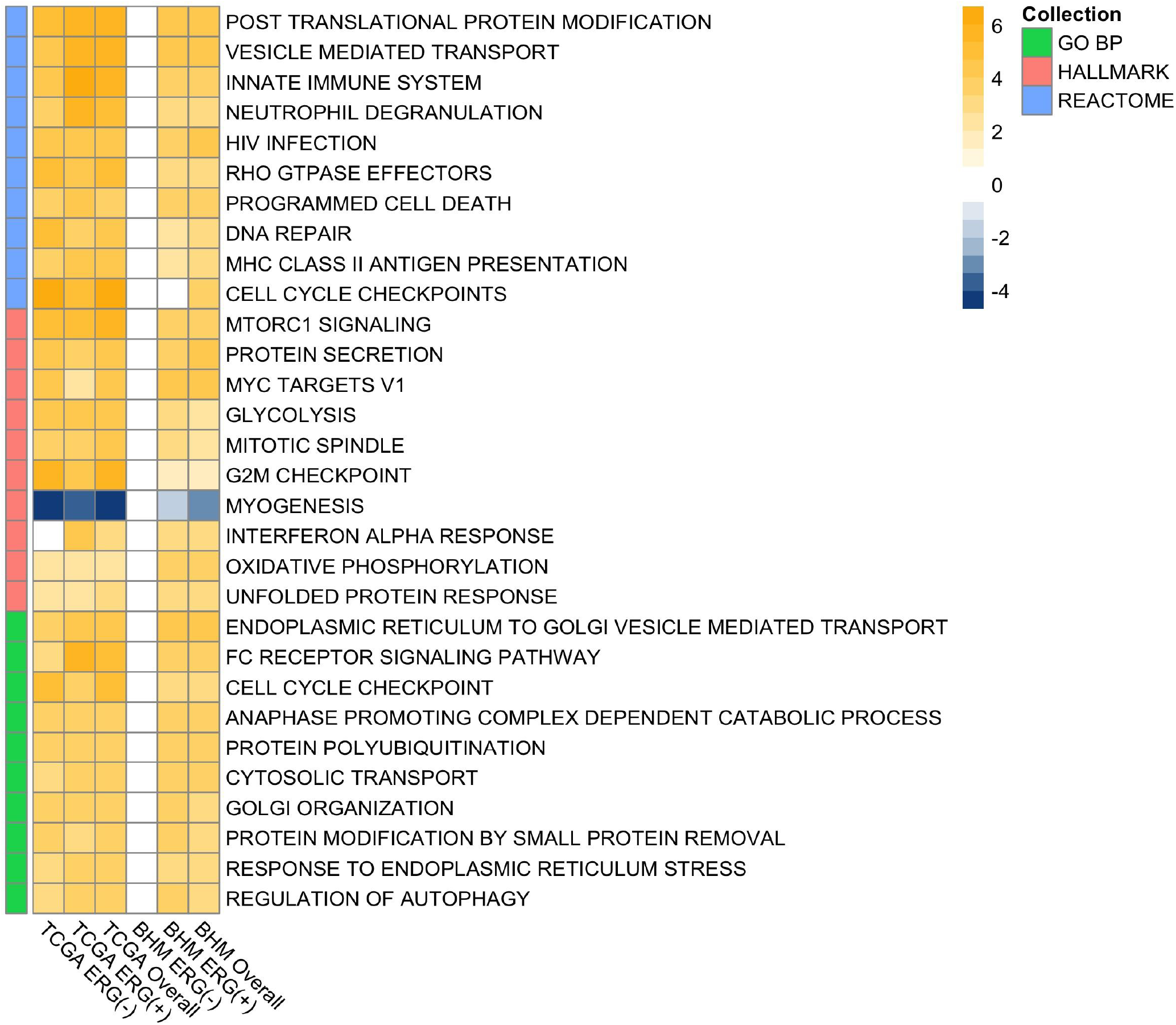
Top enriched gene sets enriched across PTEN-null and PTEN-intact in the TCGA and BHM cohorts stratified by ERG status and overall. Heatmap of mean-centered log2 signed p-values (normalized enriched score multiplied by log_10_ of p-value) showing the top 10 enriched gene sets of each collection (ranked by signed p-value).

**Figure 5.**
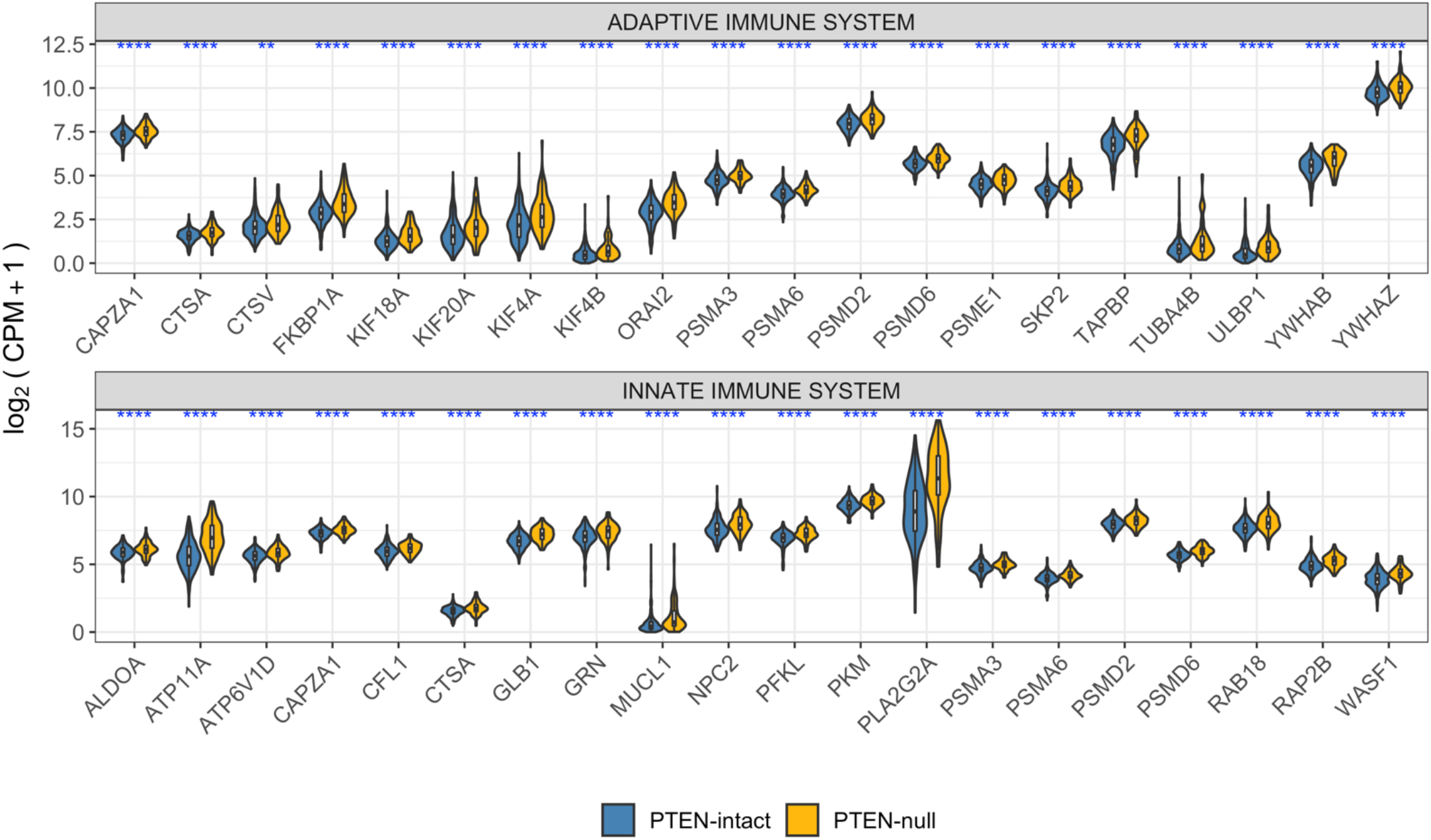
Expression of immune-related genes stratified by PTEN status. Top 20 were selected based on the leading edge of the GSEA of the adaptive and innate immune system gene sets from REACTOME. Significances based on t-test between PTEN-null and PTEN-intact using log2 CPM+1 values. Significance cutoffs: *=≤ 0.05; **≤0.01; *** ≤ 0.001; ****≤0.0001.

Since *PTEN*-null tumors are known to have decreased androgen output, which is a strong suppressor of inflammatory immune cells (29,56,57), we hypothesized that this decrease in androgen levels could activate an immune response. We, therefore, performed a GSEA analysis using a collection of androgen-regulated genes from Schaeffer et al. (29) to test if the *PTEN*-null signature was enriched in this gene set. Both the TCGA- and BHM-signature were shown to be positively enriched in genes that were shown to be repressed upon dihydrotestosterone treatment (NES =1.39-155, FDR ≤ 0.05) (Figure S8).

## Discussion

With an estimated prevalence of up to 50%, *PTEN* loss is recognized as one of the major driving events in PCa (58). *PTEN* antagonizes PI3K-AKT/PKB and is a key modulator of the AKT-mTOR signaling pathways which are important in regulating cell growth and proliferation. Accordingly, *PTEN* loss is consistently associated with more aggressive disease features and poor outcomes. Saal and collaborators previously generated a transcriptomic signature of *PTEN* loss in breast cancer (6). While this signature was correlated with worse patient outcomes in breast and other independent cancer datasets, including PCa, the signature unsurprisingly fails to capture key characteristics of PCa such as *ERG*-rearrangement (6,11). Significantly, a transcriptomic signature reflecting the landscape of *PTEN* loss in PCa has not been described to date.

Immunohistochemistry (IHC) assay is a clinically utilized technique to determine the status of the *PTEN* gene, with high sensitivity and specificity for underlying genomic deletions (59) (Figure 1). Therefore, we analyzed transcriptome data from two large PCa cohorts – the Health Professional Follow-up Study (HPFS) and the Natural History (NH) study – for which IHC-based PTEN and ERG status was available (n = 390 and 207, respectively), deriving a *PTEN*-loss gene expression signature specific to PCa (Figure 2 and Supplementary Table S1). Genes that are associated with increased proliferation and invasion in several cancer types, such as *DPT, ANPEP* and *LRRN1*, were among the most concordant DEG in this signature. *DPT* has been shown to inhibit cell proliferation through MYC repression and to be down-regulated in both oral and thyroid cancer (60,61). It has also been shown to control cell adhesion and invasiveness, with low expression leading to a worst prognosis (61,62). *ANPEP* is known to play an important role in cell motility, invasion, and metastasis progression (62,63), and lower expression of this gene has been associated with the worst prognosis (64). *LRRN1* is a direct transcriptional target of *MYCN*, and an enhancer of EGFR and IGRF signaling pathway (65). Higher levels of *LRRN1* expression promote tumor cell proliferation, inhibiting cell apoptosis, and play an important role in preserving pluripotency-related proteins through *AKT* phosphorylation (65–67), leading to a poor clinical outcome in gastric and brain cancer.

Notably, *ERG* was shown to be upregulated in our signature, which led us to perform a stratified analysis to avoid capturing signals driven mostly by *ERG* overexpression. Surprisingly, we were not able to detect significant differences by *PTEN* status in the HPFS and NH cohorts, which were quantified by gene expression microarrays, in the *ERG*^-^ samples. Conversely, when analyzing the TCGA cohort, we were able to detect significant changes by *PTEN* status in the *ERG*^-^ samples (Supplementary Tables S3-S5). However, given the known limitations of gene expression microarrays performed on formalin fixed material, such as the limited dynamic range of expression values (68), we believe that the HPFS and NH datasets were limited by the technology employed. Nevertheless, concordance between the BHM- and TCGA-cohorts were similar in both the overall and the *ERG*^+^ background comparison (Supplementary Figure S4).

We observed in the TCGA cohort several lncRNAs that have already been associated with PCa progression were found in our signature. PCA3 acts by a variety of mechanisms such as down-regulation of the oncogene *PRUNE2* and up-regulation of the *PRKD3* gene by acting as a miRNA sponge for *mir-1261* leading to increase proliferation and migration(37,38). Conversely, knockdown of *PCA3* can lead to partial reversion of epithelial-mesenchymal transition (EMT) (39) which can lead to increased cell invasion, motility, and survival (40). Although *KRTAP5-AS1* has not been associated with PCa, it has recently shown that *KRTAP5-AS1* can act as a miRNA sponge for miRNAs, such as *mir-596*, which targets the oncogene *CLDN4* which enhances the invasion capacity of cancer cells and promote EMT (40,43), thereby overexpression of *KRTAP5-AS1* can lead increased levels of *CLDN4* (44). *Mir-596* has also been shown to be overexpressed in response to androgen signaling and associated with anti-androgen therapy resistance (44).

Moreover, many lncRNAs exclusively annotated in the FANTOM-CAT were associated with PTEN-loss and were shown to be expressed mostly in PCa (Figure 3). Since these genes are novel genes without elucidated function, we analyzed potential roles for these genes by looking at coding genes located in the same loci. Among the top DE lncRNAs, genes within proximity to coding genes were *CATG00000038715, CATG00000079217*, and *CATG00000117664* (Figure S6) which are positioned in the same loci as *CYP4F2, FBXL7*, and *GPR158*, respectively. *CYP4F2* is involved in the process of inactivating and degrading leukotriene B4 (*LTB4*). *LTB4* is a key gene in the inflammatory response that is produced in leukocytes in response to inflammatory mediators and can induce the adhesion and activation of leukocytes on the endothelium.(69). *FBXL7* regulates mitotic arrest by degradation of *AURKA*, which is known to promote inflammatory response and activation of NF-κB (70,71). Likewise, increase expression of *GPR158* is reported to stimulate cell proliferation in PCa cell lines, and it is linked to neuroendocrine differentiation (72).

We consistently observed a strong enrichment in immune response genes and gene sets upon *PTEN* loss (Figure 4 and Supplementary Tables S6-S20). Immune-associated genes (i.e. *GP2* and *PLA2G2A)* were found amongst the top up-regulated genes in our signature (Figure 2). Positive enrichment of Interferon-alpha- and gamma-response genes (FDR ≤ 0.01) further suggests that a strong immuno-responsive environment, with both innate and adaptive systems activated, is developed in *PTEN*-null tumors (Figure 5). The positive enrichment of MHC class II antigen presentation, neutrophil degranulation, vesicle-mediated transport, and FC receptor pathway-related genes suggests that *PTEN*-null tumors may be immunogenic (Figure 4). This finding was particularly surprising given that *PTEN* is itself a key positive regulator of innate immune response, controlling the import of *IRF3*, which is responsible for IFN production. Accordingly, disruption of PTEN expression has previously been reported to lead to decreased innate immune response (73). Conversely, it has also been hypothesized that the increased genomic instability caused by, or associated with, *PTEN* loss can increase immunogenicity in the tumor micro-enviroment (TME) (74). This finding is of particular interest given that immune-responsive tumors can be good candidates for immunotherapy-based approaches.

Remarkably, despite loss of *PTEN* being associated with higher expression of the immune checkpoint gene programmed death ligand-1 (*PD-L1*) in several cancer types (75,76) this is not true in PCa (77). So far, current immunotherapeutic interventions, such as *PD-1* blockade, in PCa have not been successful. One of the possible reasons is the lack of *PD-L1* expression (77). Therefore, alternative targets must be considered for immunotherapy in PCa. One alternative target is the checkpoint molecule *B7-H3* (*CD276*), whose expression has already been associated with PCa progression and worse prognosis (78) and has been suggested as a target for immunotherapy (79,80). *CD276* was one of the most concordant up-regulated genes in our signature (Figure 2) suggesting that its expression is associated with *PTEN* loss. Interestingly, *B7-H3* expression may be down-regulated by androgens (81).

The effects of androgen on the immune system has already been extensively studied and reviewed (56). Androgens are known to suppress inflammatory immune cells and to impair the development and function of B- and T-cells (57). We, therefore, hypothesized that the decreased levels of androgen in *PTEN*-null TME could lead to an unsuppressed immune system. By testing our signature for enrichment in androgen-related genes (AR) derived from Schaeffer et al. (29), we observed that upon *PTEN*-loss, androgen-sensitive genes that are typically suppressed by DHT are positively enriched, indicating that androgen levels or androgen response in *PTEN*-null tumors may be lower than in their *PTEN*-intact counterparts (Figure S8). This decrease in AR-signaling has been described in *PTEN*-null tumors, in which activation of PI3K pathway inhibits AR activity. (82). Furthermore, AR inhibition activates AKT signaling by inhibiting AKT phosphatase levels further boosting cell proliferation (82), which has also been noted in this study (Figure 3). Finally, in the non-coding repertoire, both *PCA3* and *PCGEM1* are modulated by androgen (83,84) and were down-regulated upon *PTEN* loss which tracks with the observed decreased androgen response in *PTEN*-null tumors (Figure S6 and S8).

## Conclusion

Altogether, we have generated a highly concordant gene signature for *PTEN* loss in PCa across three independent datasets. We show that this signature was highly enriched in proliferation and cell cycle genes, leading to a more aggressive phenotype upon *PTEN* loss, which is concordant with the literature. Moreover, we have shown that *PTEN* loss is associated with an increase in both innate and adaptive immune response. Although the literature shows that *PTEN* loss usually leads to immuno-suppression, we find evidence that this finding may be reversed in PCa. This observation has potential implications in the context of precision medicine since immune responsive tumors are more likely to respond to immunotherapies. Therefore, *PTEN*-null tumors might benefit more from this approach than *PTEN-* intact tumors. Potentially, *PTEN* status can guide immunotherapy combination with other approaches such as androgen ablation.

Finally, by leveraging the FC-R2 resource, we were able to highlight many lncRNAs that may be associated with PCa progression. Although functional characterization these lncRNAs is beyond the scope of this study, we have shown that these novel lncRNAs are highly specific to PCa and track with several coding mRNAs and lncRNAs already reported to be involved in PCa development and progression, most notably, genes involved in immune response. By providing a PCa-specific signature for PTEN loss, as well as highlighting potential new players, we hope to empower further studies on the mechanisms leading to the development of PCa as well its more aggressive subtypes aiding in the future development of potential biomarkers, drug targets and guide therapies choice.

## Acknowledgments

This publication was made possible through support from the National Institutes of Health–National Cancer Institute (NIH-NCI) grants U01CA196390, and R01CA200859; and the U.S. Department of Defense (DoD) office of the Congressionally Directed Medical Research Programs (CDMRP) award W81XWH-16-1-0739 and W81XWH-16-1-0737; and Fundação de Amparo a Pesquisa do Estado de Minas Gerais award BDS-00493-16.

## Author contributions statement

L.M. and T.L. conceived the idea; L.M., E.L.I. and T.L. designed the study; E.L.I., D.F.S., W.D., T.L., and L.M. performed the analysis; E.L.I., D.F.S., T.L., T.V., G.R.F., and L.M. interpreted the results; T.L., E.M.E., S.T., L.M., M.L., and E.M.S. provided data and tools; E.L.I., D.F.S., T.L., and L.M. wrote the manuscript; all authors reviewed and approved the manuscript.

## Supplementary Figures and Tables

**Figure S1.**
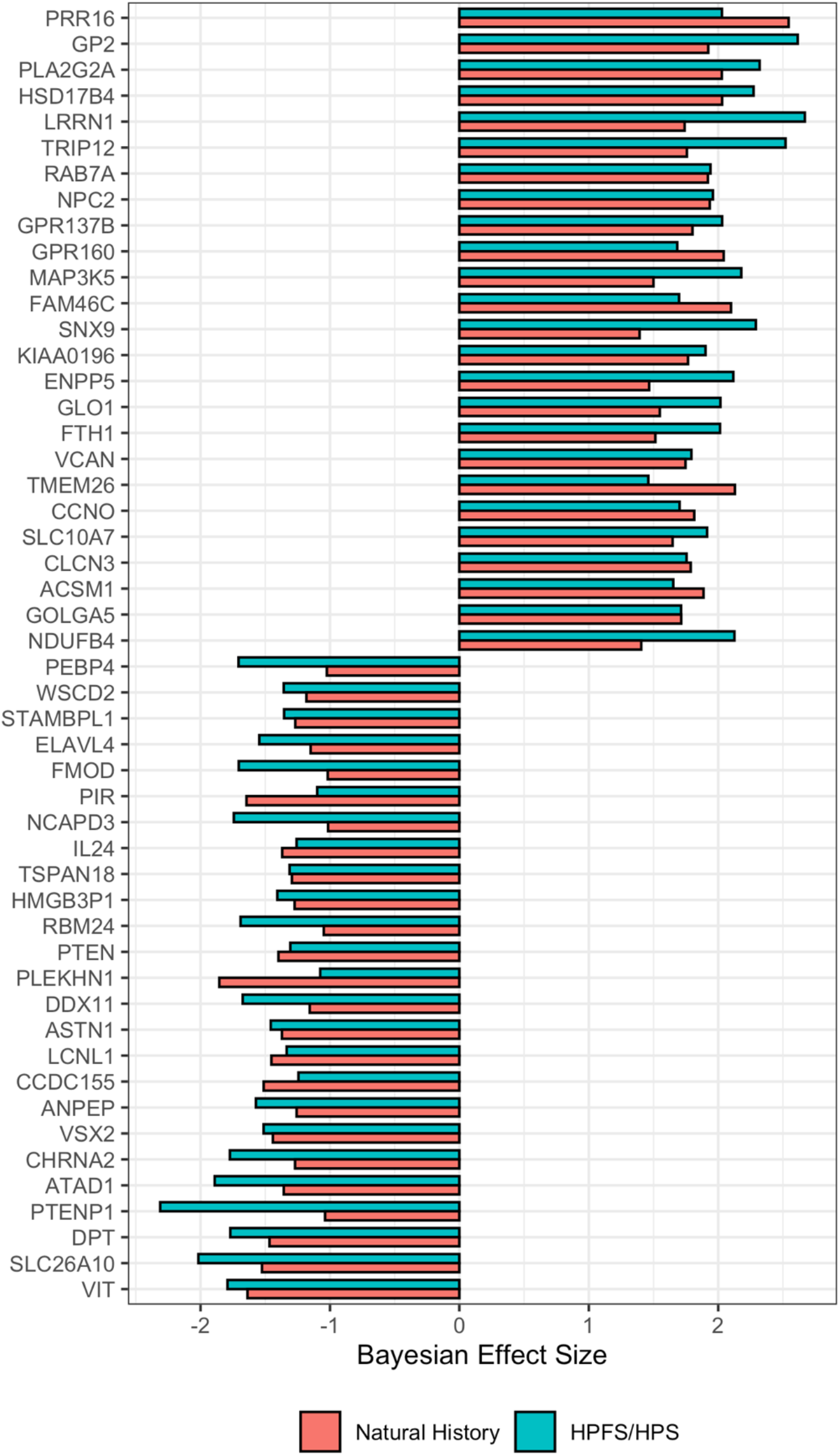
Cross-study of differential gene expression in PTEN-null vs PTEN-intact in ERG^+^ samples. Meta-analysis of HPFS/PHS and NH cohorts with Bayesian Hierarchical Model for DGE using XDE showing the top 25 most concordant differentially up- and down-regulated genes. PTEN status were based on IHC assays.

**Figure S3.**
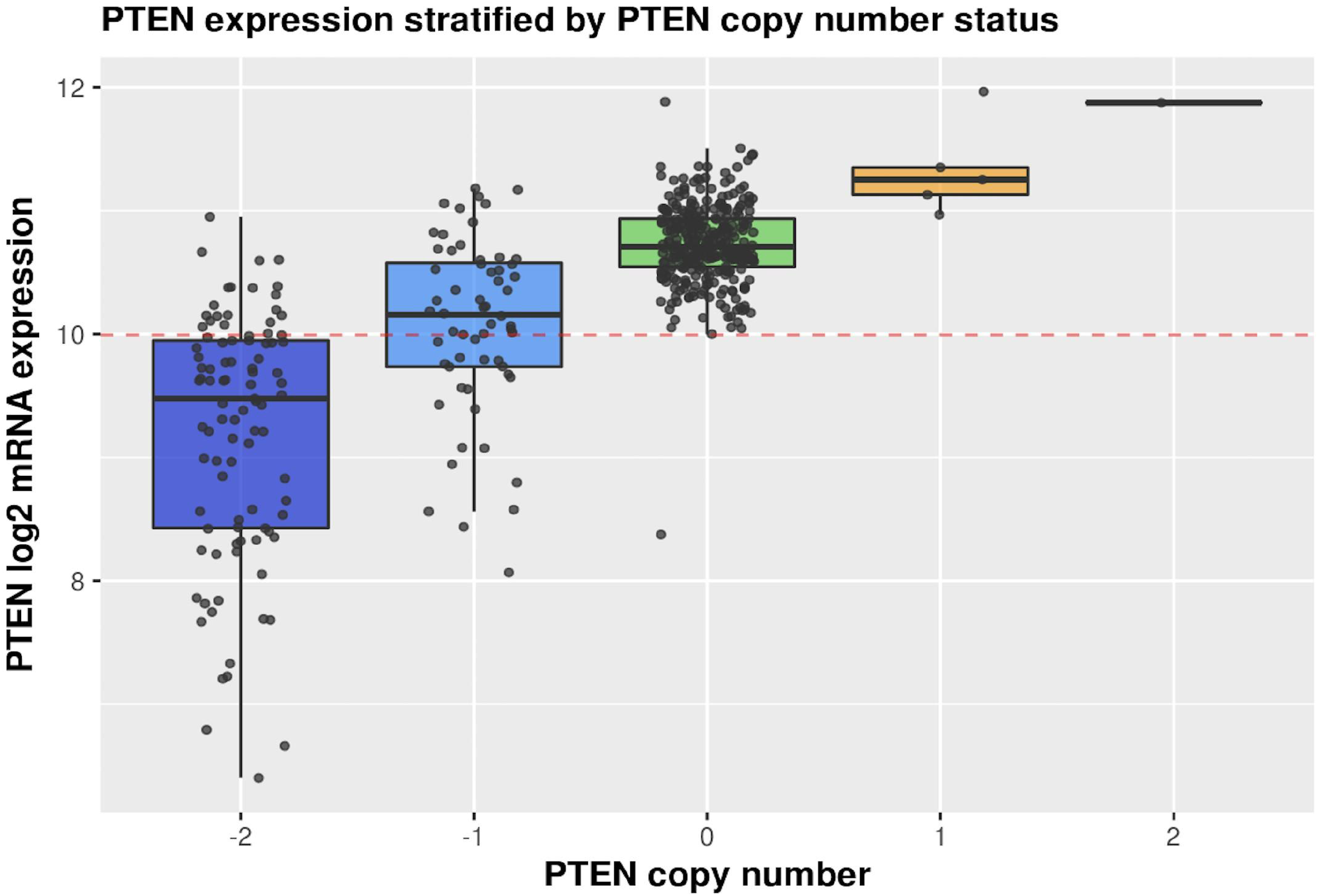
PTEN expression levels stratified by CNV. Figure shows PTEN expression levels distribution by copy number variation (CNV), called by GISTIC algorithm.

**Figure S4.**
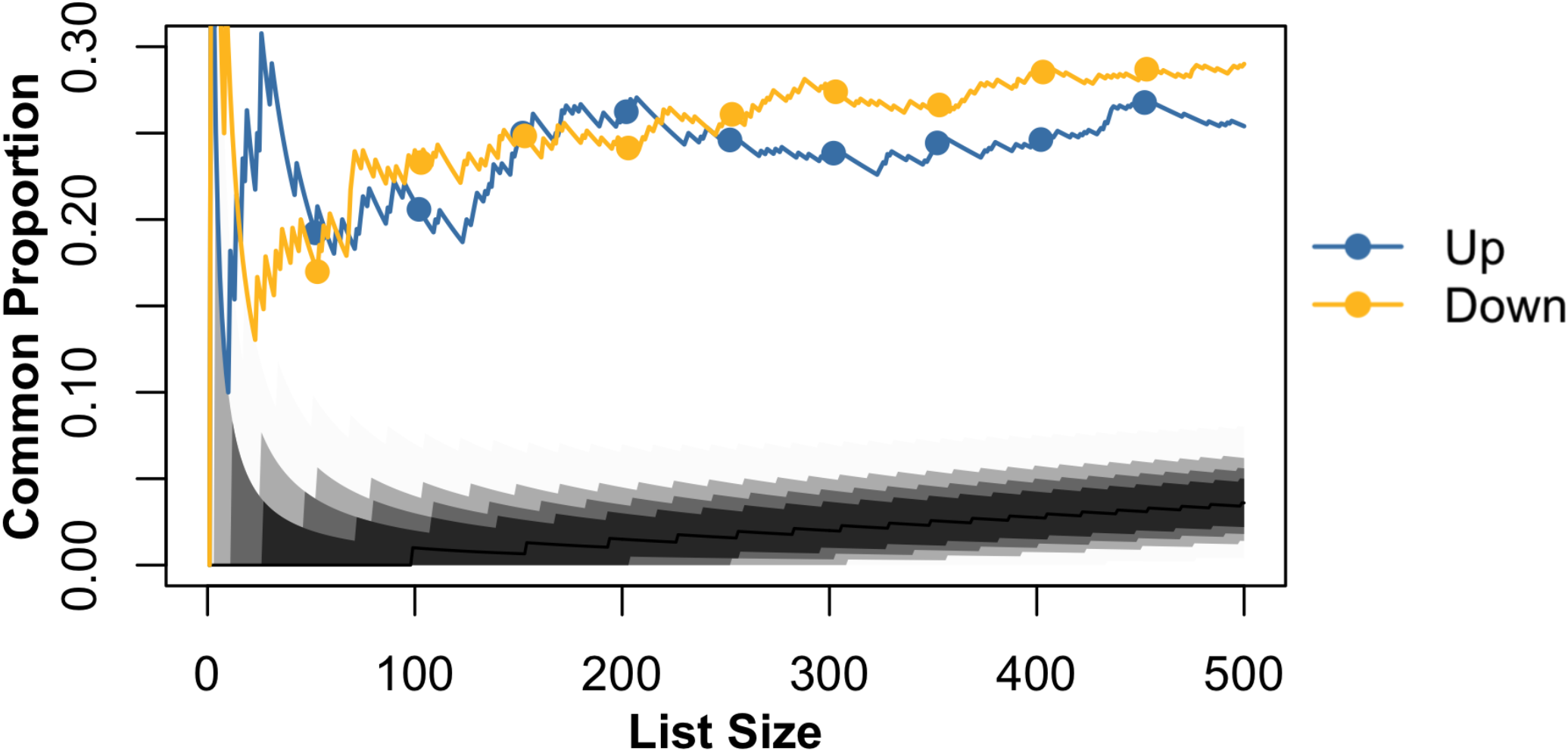
Correspondence-at-the-top (CAT) plot between TCGA CNV-based calls and the Bayesian Hierarchical Model approach (BHM). Agreement of genes ranked by t-statistics (TCGA) and average Bayesian Effect Size (BHM). Lines represent agreement between tested cohorts for PTEN-intact vs PTEN-null. Black-to-light grey shades represent the decreasing probability of agreeing by chance based on the hypergeometric distribution, with intervals ranging from 0.999999 (light grey) to 0.95 (dark grey). Lines outside this range represent agreement in different cohorts with a higher agreement than expected by chance.

**Figure S5.**
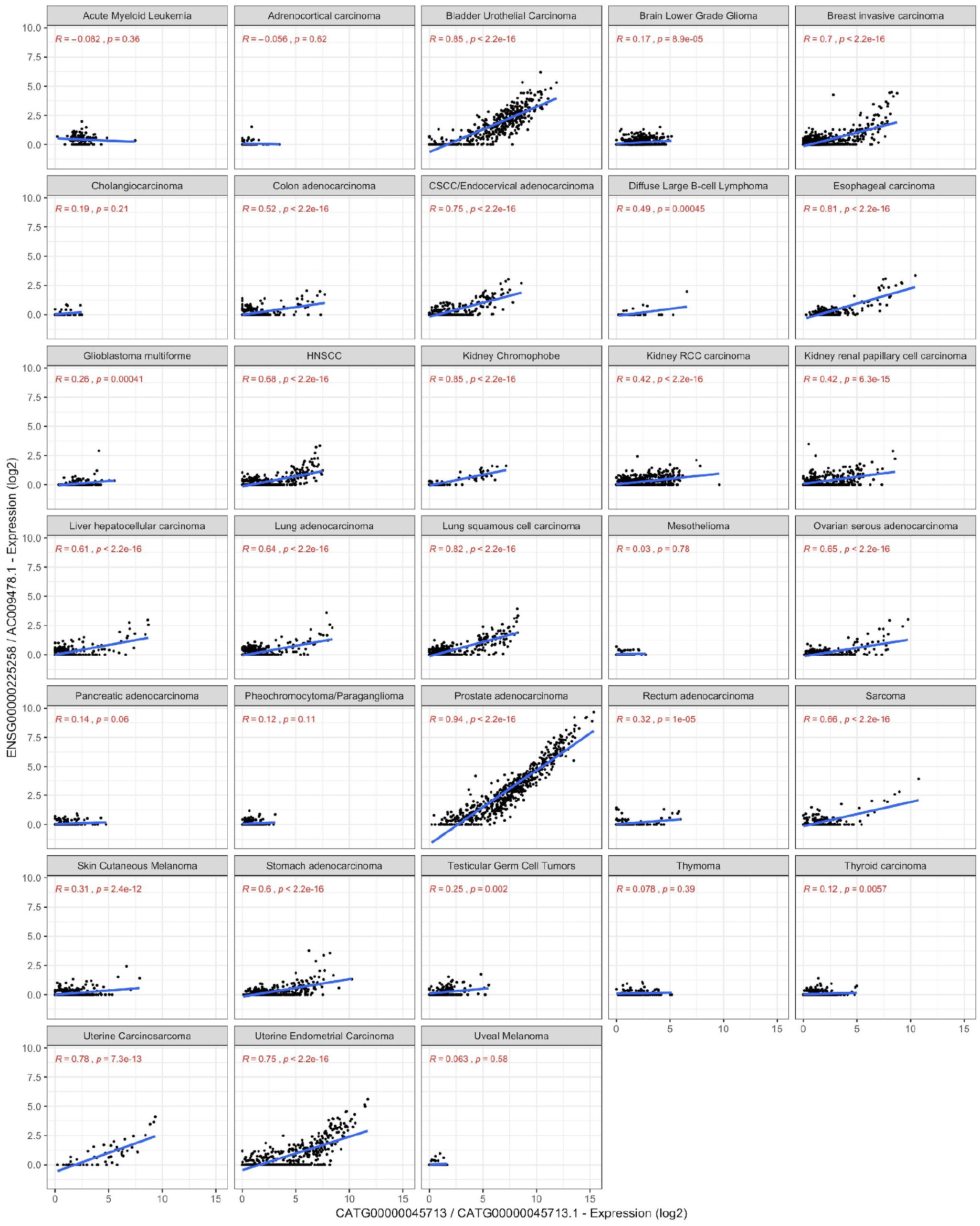
Expression of AC009478.1 is shown to be highly specific to PRAD, BLCA, to a lesser extent in UECA and BRCA. Figure shows raw expression values of SchLAP1 and AC009478.1 across cancer types. Pearson correlations and p-values are shown in red.

**Figure S6.**
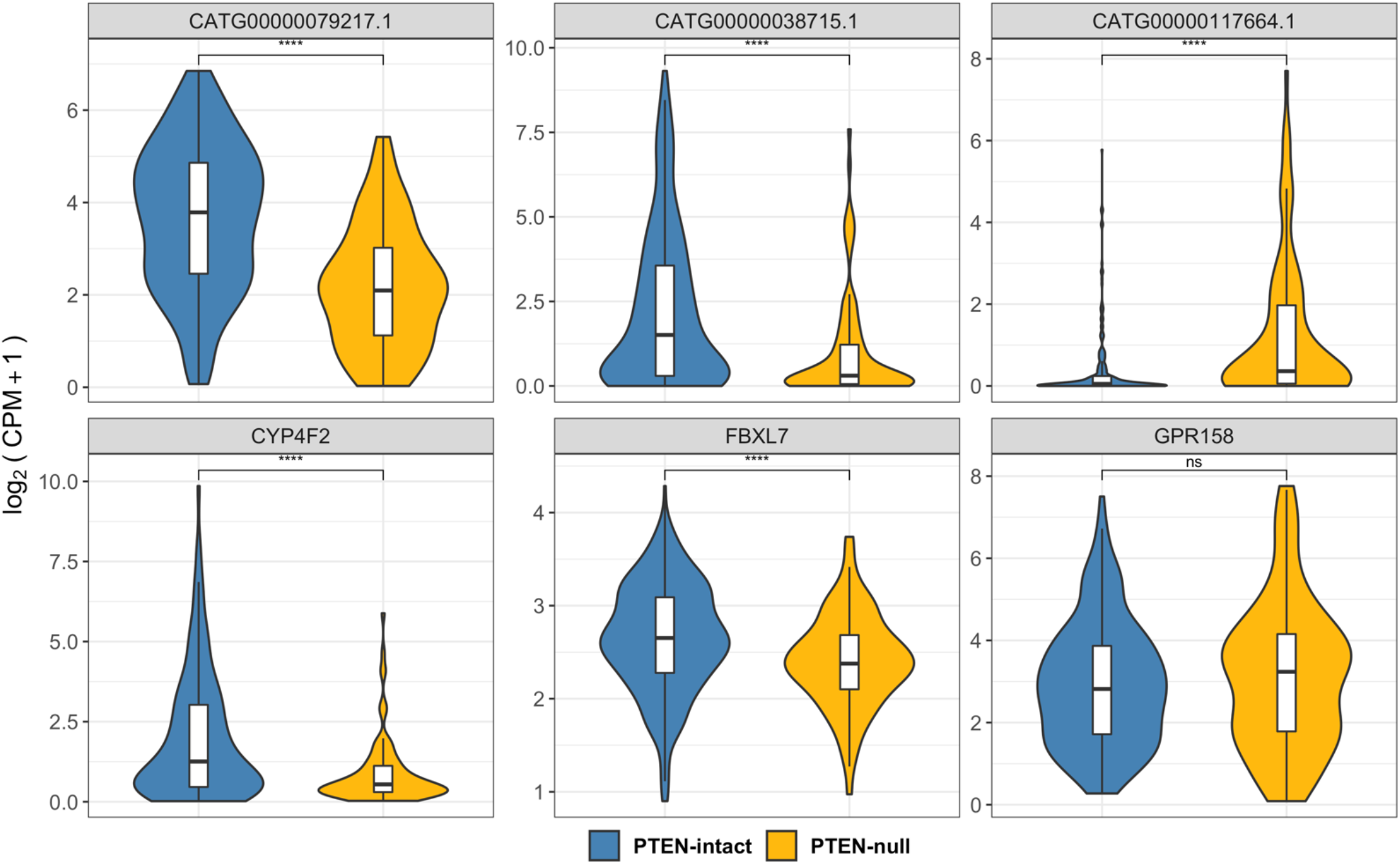
Expression of FANTOM-CAT lncRNAs genes (top) and close coding genes (bottom) stratified by PTEN status. Significances based on t-test between PTEN-null and PTEN-intact using log2 CPM+1 value. Significance cutoffs: *=≤ 0.05; **≤0.01; *** ≤ 0.001; ****≤0.0001.

**Figure S7.**
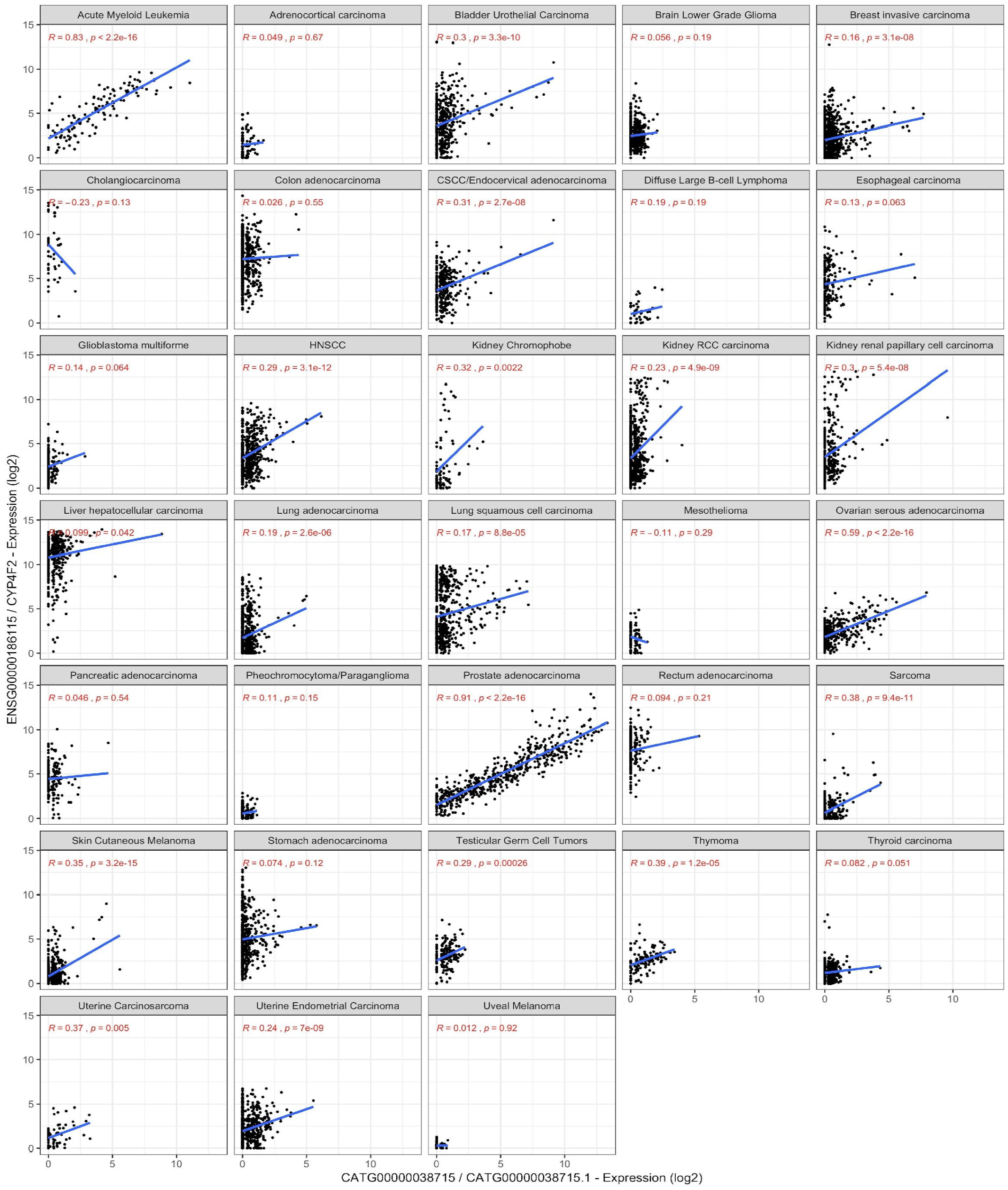
Person correlation gene CATG00000038715 and CYP4F2 across cancer types. CATG00000038715 and CYP4F2 expression are shown to be highly correlated in PCa. Moreover, CATG00000038715 expression is shown to be highly specific to PCa. With exception of leukemia cells, none of the other tumors expressed high levels of CATG00000038715.

**Figure S8.**
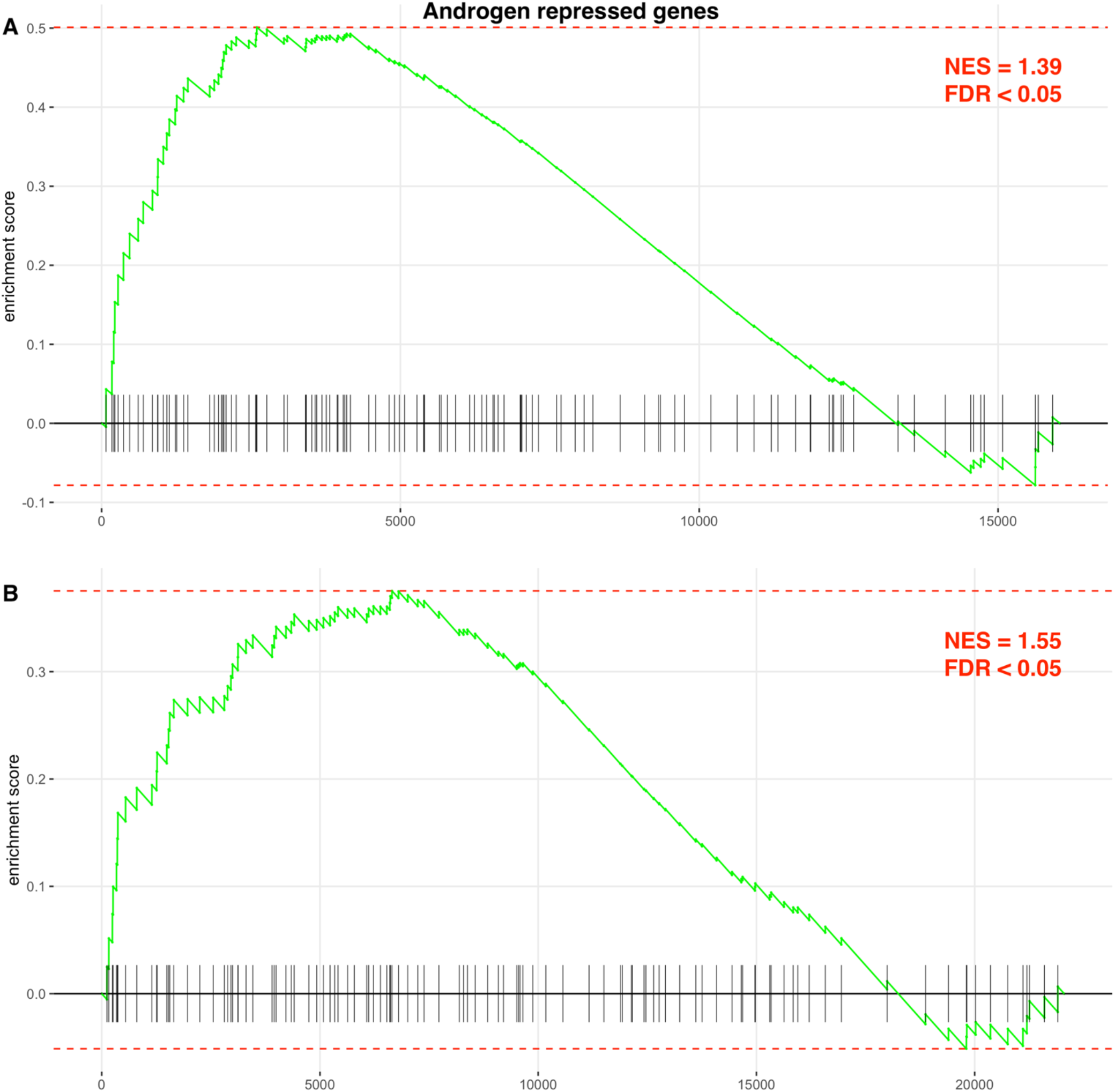
Gene set enrichment for Androgen repressed genes. Gene set enrichment analysis of gene signature showing positive enrichment of genes repressed by dihydrotestosterone after 6 hours of exposure obtained from Schaeffer et al.^48^. Enrichment for BHM-signature is shown in panel A and TCGA-signature in panel B.

